# Machine-Perception Nanosensor Platform to Detect Cancer Biomarkers

**DOI:** 10.1101/2021.04.28.441499

**Authors:** Zvi Yaari, Yoona Yang, Elana Apfelbaum, Alex Settle, Quinlan Cullen, Winson Cai, Kara Long Roche, Douglas A. Levine, Martin Fleisher, Lakshmi Ramanathan, Ming Zheng, Anand Jagota, Daniel A. Heller

## Abstract

Conventional molecular recognition elements, such as antibodies, present issues for the development of biomolecular assays for use in point-of-care devices, implantable/wearables, and under-resourced settings. Additionally, antibody development and use, especially for highly multiplexed applications, can be slow and costly. We developed a perception-based platform based on an optical nanosensor array that leverages machine learning algorithms to detect multiple protein biomarkers in biofluids. We demonstrated this platform in gynecologic cancers, which are often diagnosed at advanced stages, leading to low survival rates. We investigated the platform for detection in uterine lavage samples, which are enriched with cancer biomarkers compared to blood. We found that the method enables the simultaneous detection of multiple biomarkers in patient samples, with F1-scores of ~0.95 in uterine lavage samples from cancer patients. This work demonstrates the potential of perception-based systems for the development of multiplexed sensors of disease biomarkers without the need for specific molecular recognition elements.

## Introduction

Current biomolecular identification methodologies rely heavily on one-to-one recognition via specific proteins and nucleic acids such as antibodies, peptides, and aptamers to bind analytes.^1–5^ However, the development of highly sensitive and specific binding moieties in a quantity sufficient to detect target molecules with one-to-one recognition has multiple challenges that delay the development of a robust, versatile, and cost-effective platform for multiple analyte detection. The challenges of using antibodies include long-term stability/robustness, transient/real-time applications, and production difficulties, especially when many different antibodies must be developed.^6–8^ Hence, technologies that replace antibodies could enhance the development of certain types of point-of-care assays, medical devices such as wearable sensors, and aid diagnostics in under-resourced settings, where cold chain storage is limited.^9,10^

Perception-based machine learning platforms, modeled after the complex olfactory system, can isolate individual signals through an array of relatively non-specific receptors.^11^ Each receptor captures certain features and the overall ensemble response is analyzed by the neural network in our brain, resulting in perception. Biofluids such as blood, urine, saliva, and sweat are indicative of physiological conditions and enable biomarker detection in their native state.^12,13^ Recent advances in machine learning (ML) methodologies have made complex algorithms more accessible, facilitating the integration of perception systems into materials science.^14,15^ We believe that perception-based sensors can be developed to enable the successful, multiplexed detection of analytes without the need for antibodies.

Previous attempts to develop perception-based sensing platforms have had limited success. Prior works include the “electronic nose” ^16–19^ for gas sensing based on conducting polymers, DNA-decorated field-effect transistors,^20^, and protein recognition using simple data analytic techniques.^21^ “Optical” noses have been developed as well.^22,23^ However, these developments are limited in their ability to detect molecules such as proteins and in physiological conditions and complex biofluids. To overcome limitations associated with one-to-one recognition elements, we are investigating the development of a perception-based methodology that uses ML processes coupled with a sensor array, where each element exhibits moderate selectivity for a wide range of molecules.

The prognosis and quality of life of cancer patients are significantly affected by the ability to accurately diagnose diseases at an early stage. One such example is ovarian cancer (OC), the fifth-leading cause of cancer-related deaths among women in the United States and first among gynecologic malignancies,^24^ with 22,000 new cases and 14,000 deaths per year.^24^ The five-year relative survival rate for patients diagnosed with ovarian cancer is 44%,^25^ while detection at stage I can increase the five-year survival rate to more than 90%.^26^ However, there are no methods to date that achieve early, accurate diagnoses, nor are there strategies to rapidly determine patient response to treatment in order to inform the choice of therapy.

To detect gynecologic cancers, such as high-grade serous ovarian carcinoma (HGSOC)^27–29^, and endometrial cancers,^30,31^ FDA-approved serum biomarkers such as cancer antigen 125 (CA-125) and human epididymis protein 4 (HE4) have been used as well as ultrasonography. However, these methods lack the sensitivity to detect early-stage cancer and have had little impact on survival.^32,33^ A recent study of uterine lavage (or uterine washings, fluids removed from the uterus after perfusion with saline) discovered significantly higher levels of biomarkers, such as HE4, CA-125, chitinase-3-like protein 1 (YKL-40), and mesothelin (MSN) than those found in serum.^34^ Therefore, the use of uterine lavage has the potential to improve early detection.

Single-wall carbon nanotubes (SWCNTs) have unique optical properties and sensitivity which make them valuable as sensor materials.^35^ SWCNTs emit near-infrared (NIR) photoluminescence with distinct narrow emission bands that are exquisitely sensitive to the local environment.^36^ In addition, the emission is photostable, enabling quantitative and long-term monitoring of small-molecules, proteins, nucleic acids, and enzymatic activities both *in vitro* and *in vivo*.^37–41^ Individual SWCNT species (or chiralities) have distinct bandgaps, which contribute to their varying sensitivities to the redox and charge phenomena.^42–44^ Coatings such as DNA can confer not only colloidal stability in an aqueous solution but also selectivity by modulating the surface coverage and bandgaps.^45^ The use of DNA-wrapped SWCNTs (DNA-SWCNTs) has been used for the detection of a wide range of analytes in biological media, including in live cells and animals.^41,46^

In this study, we investigate a machine-perception (MP)-based sensing system to detect multiple biomarkers in human biofluids (Fig. 1). We developed a DNA-SWCNT-based photoluminescent sensor array wherein the optical responses were used to train machine learning models to detect gynecologic cancer biomarkers HE4, CA-125, and YKL-40 in laboratory-generated samples and patient fluids. Distinct changes in fluorescent peak position and intensity values from each DNA-SWCNT combination were observed in response to the protein analytes. Machine learning algorithms support vector machine (SVM), random forest (RF), and artificial neural network (ANN) enabled the prediction of both the presence (classification) and concentration (regression) of each biomarker. In uterine lavage samples, the classification results were highly accurate, producing F1-scores of ~0.95 in laboratory-generated samples and classification successes of 100% for HE4 and CA-125 and 91% for YKL-40 in cancer patient samples. This work suggests that a nanosensor/perception-based sensing system can accurately detect multiple disease biomarkers in patient biofluids.

**Figure 1.**
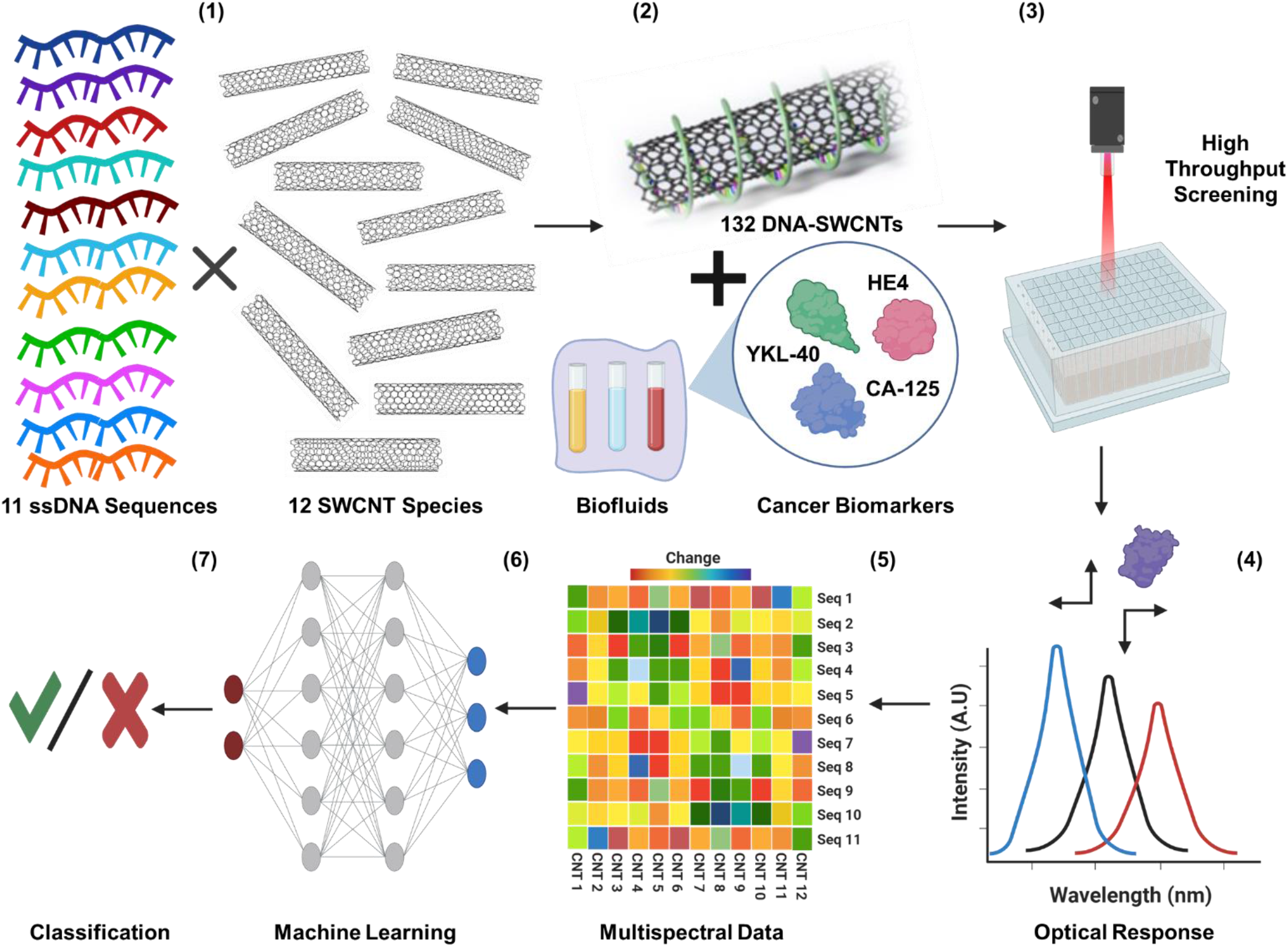
Machine-Perception Nanosensor Platform. **(1)** Eleven single-stranded DNA oligonucleotides wrap SWCNT chiralities to form DNA-SWCNT sensor complexes. **(2)** The array of sensors is incubated in the sample of interest. **(3)** The optical response of the sensors is interrogated by high-throughput near-infrared spectroscopy. **(4)** The spectroscopic data are fitted to determine the wavelength and intensity of each sensor emission band. **(5)** The sensor responses are processed into a feature vector training set. **(6)** Machine learning algorithms are trained and validated for each target protein and their combinations. **(7)** Prediction results are evaluated.

## Results

### DNA-SWCNT array

We characterized multiple DNA-SWCNT complexes to form the basis of a sensor array. Eleven DNA sequences ((AT)_11_, (AT)_15_, (AT)_20_, (GT)_12_, (ATT)_4_, (TCT)_5_, T_3_C_3_T_3_C_3_T_3_, C_3_T_9_C_3_,C_3_T_3_C_9_,CT_2_C_3_T_2_C, and (AC)_15_) were chosen because many of them are recognition sequences of specific SWCNT chiralities, which suggest ordered wrapping on their surface, while others confer some degree of specificity to proteins or other analytes.^47–50^ Twelve semiconducting SWCNT species present in the HiPCO preparation ((6,5), (8,4), (10,3), (7,5), (7,6), (8,3), (9,5), (9,4), (8,6), (8,7), (10,2), and (9,7)) were evaluated due to their high concentrations in the sample and bright photoluminescence in the serum/water optical window of 900-1400 nm (Fig. S1A, B). The combinatorial possibilities of 12 SWCNT species and 11 DNA sequences resulted in the formation of 132 distinct DNA-SWCNT complexes that were investigated within the context of a sensor array. The DNA-SWCNT complexes exhibited high colloidal stability and strong photoluminescence, as previously reported.^51–53^ We characterized the complexes using UV-Vis-NIR absorbance, NIR fluorescence spectroscopy, atomic force microscopy, and zeta potential measurements (Fig. S1). The measurements confirmed the emissive properties of at least 12 SWCNT chiralities (Fig. S1B), a highly negative zeta potential of DNA-SWCNT complexes formed with all 11 DNA sequences (Fig. S1C), and a DNA banding pattern along the SWCNT surface for all sequences (Fig. S1D-H).

The optical responses of the DNA-SWCNT complexes to known gynecologic cancer biomarkers were assessed via spectroscopy. High-throughput, near-infrared spectroscopy (in the range of 900-1400 nm) was conducted on all DNA-SWCNT complexes introduced to lab-generated samples of the protein biomarkers HE4, CA-125, and YKL-40 in 10% FBS solutions (to provide a relevant background of interferent molecules). The spectroscopic bands of all SWCNT chiralities were fitted to extract peak wavelength shift (Δλ) and intensity ratio (I/I_0_) with respect to a control sample in 10% FBS. As a representative example, the (7,5) chirality emission peak blue-shifted (Δλ<0) and its intensity was attenuated (I/I_0_ <1) in response to HE4 (Fig. 2A), while brightening and red-shifting were observed upon exposure to CA-125 and YKL-40 (Fig. 2A and inset). Similar analyses found diverse optical responses to single biomarkers across SWCNT chiralities (Fig. 2B, C) and DNA wrappings (Fig. 2D, E). There were no obvious correlations between the response and conditions in which they were challenged (Fig. 2F, G, and Fig. S2).

**Figure 2.**
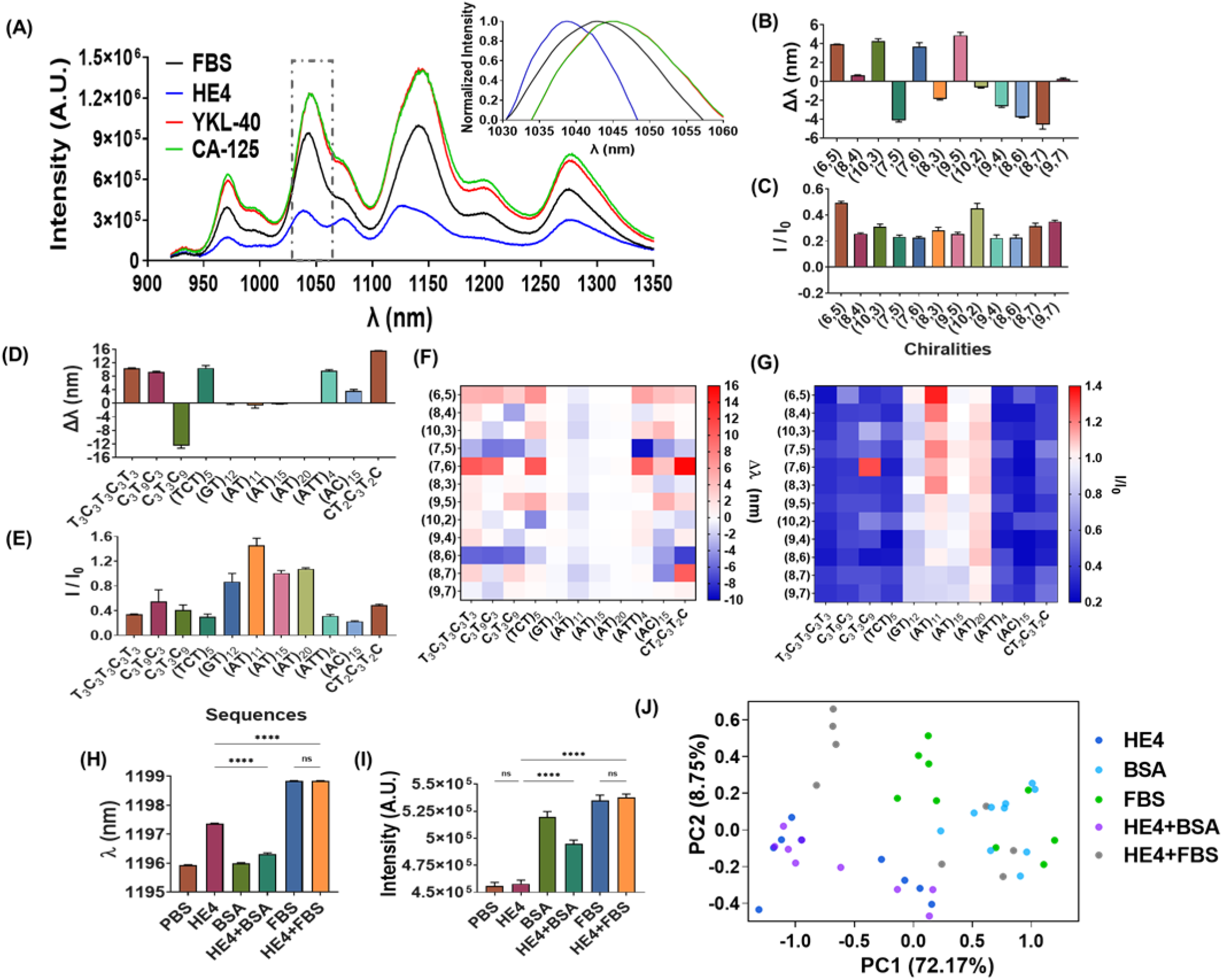
DNA-SWCNT optical responses to gynecologic cancer biomarkers. **(A)** Representative spectra of DNA-SWCNT complexes in response to cancer protein biomarkers. Inset: normalized spectrum of the (7,5) chirality. **(B)** Wavelength modulation of (AC)_15_-SWCNT complexes upon incubation with 100 nM of HE4; n=3. **(C)** Intensity modulation of (AC)_15_-SWCNT complexes upon incubation with HE4; n=3. **(D)** Wavelength modulation of DNA-(7,6) complexes upon incubation with HE4; n=3. **(E)** Intensity modulation of DNA-(7,6) complexes upon incubation with HE4; n=3. **(F)** Heatmap of total wavelength modulations of DNA-SWCNT complexes upon incubation with HE4; n=3. **(G)** Heatmap of total intensity modulations of DNA-SWCNT complexes upon incubation with HE4; n=3. **(H)** Wavelength of (AT)_11_-(8,6) complex upon incubation with PBS, HE4, BSA, and FBS in PBS; n=3, mean ± SEM; ****P < 0.0001, unpaired t-test. **(I)** Intensity of (AT)_11_-(8,6) complex upon incubation with PBS, HE4, BSA, FBS in PBS; n=3, mean ± SEM; ****P < 0.0001, unpaired t-test, “ns” denotes not significant. **(J)** PCA plot of the DNA-SWCNT response to HE4 versus interferents.

To study the physical properties of the DNA-SWCNT complexes that could contribute to the distinct responses, we analyzed the SWCNT surface charge and DNA wrapping patterns on the SWCNT surface. Zeta potential measurements of the DNA-SWCNTs showed that surface charge varied between approximately −44 and −55 mV, depending on the DNA sequence (Fig. S1C), likely a result of differences in DNA packing densities. To further investigate, we conducted atomic force microscopy (AFM), which revealed significant differences in the density of observable height maxima/peaks on the SWCNTs of approximately 40% (Fig. S1D-H), ascribable to the DNA. These findings suggest that the unique responses of each DNA-SWCNT to the proteins are likely due in part to the distinct DNA wrapping patterns on each SWCNT chirality.

Next, we investigated the specificity of the DNA-SWCNTs by examining the response to HE4 in the presence of interferents (i.e. bovine serum albumin (BSA) and fetal bovine serum (FBS)). We found that some DNA-SWCNTs responded differently to the analyte and interferents, but the specificity of any one complex appeared marginal (Fig. 2H, I). To assess the distinctness of DNA-SWCNT responses to a protein biomarker vs. interferents, we applied principal component analysis (PCA). The analysis of DNA-SWCNT responses to HE4 and interferent proteins failed to separate distinct optical responses of HE4 (Fig. 2J). We thus concluded that more sophisticated data analyses were needed to determine whether the DNA-SWCNT array could correctly identify analytes within a complex environment.

### ML feature vector construction

In order to differentiate the biomarkers via the DNA-SWCNT optical responses, we investigated several machine learning strategies. We tested two different feature vector (FV) methods to represent experimentally measured data matrices composed of DNA sequences and SWCNT chiralities (Fig. 3A). Each vector was constructed with two components: “*Example ID*” – SWCNT chirality or DNA sequence, and *“Features” –* the DNA-SWCNT complex emission intensity and wavelength response. In addition, the vector corresponds to a specific label that indicates the presence of each biomarker in the sample. The first feature vector (FV_1_) is focused on chirality (Fig. 3A_(1)_) and uses DNA sequences as the example IDs and chirality-dependent optical responses as features. Underlying this choice of feature is the hypothesis that the spectroscopic response of multiple SWCNTs in combination with a single DNA sequence is sufficient to determine the presence or concentration of biomarkers. DNA sequences were encoded into IDs as either bi-gram or tri-gram term frequency vectors.^47^ Therefore, the total number of features is 40 using a bi-gram representation (16+2*12) and 88 using a tri-gram frequency vector (64+2*12).

**Figure 3.**
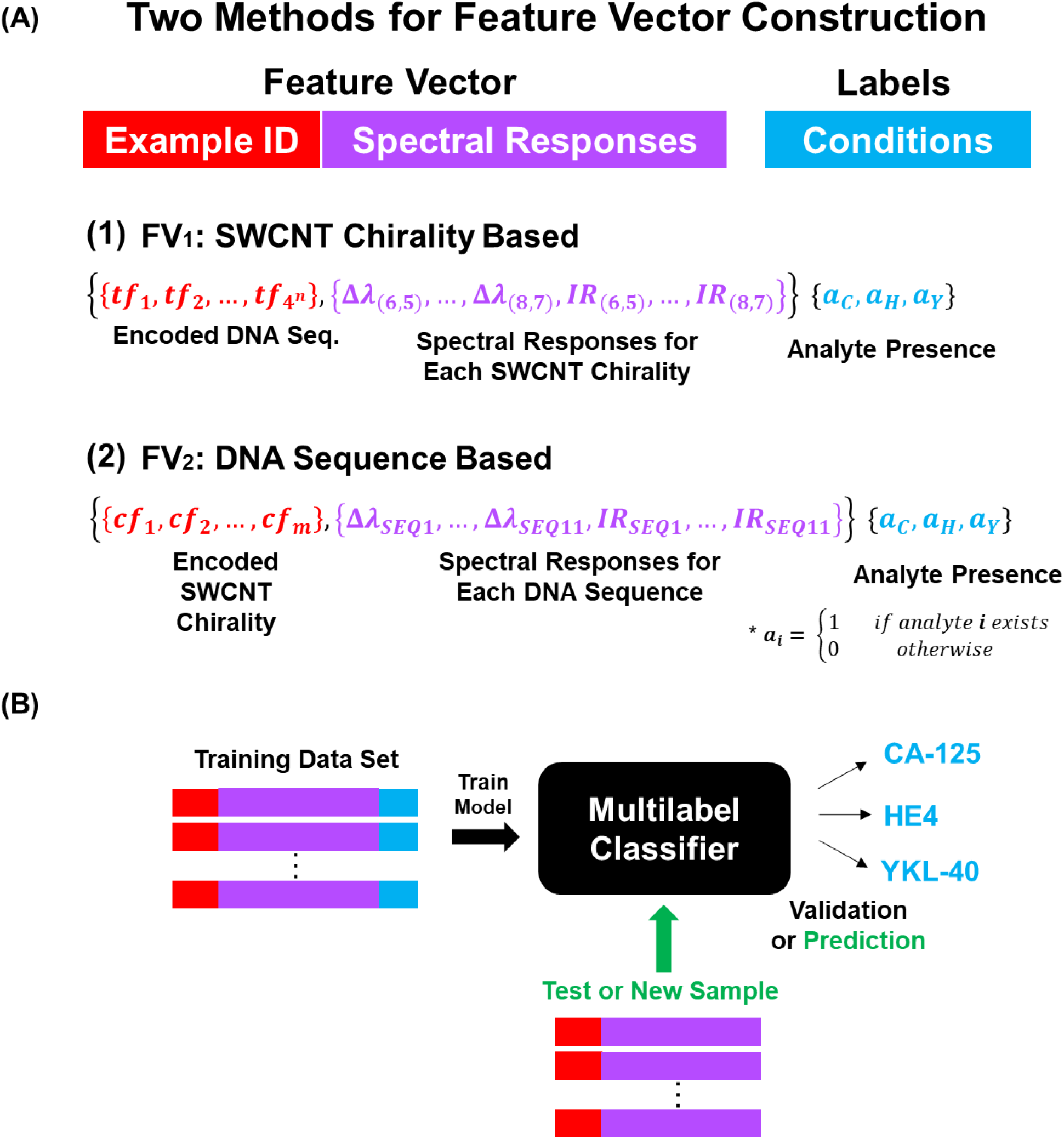
Feature vector construction. **(A)** The feature vector contains two parts – example encoding (red) and optical response-based features (purple) – with each vector corresponding to a label that indicates the biomarker conditions (blue). The total features of FV_1_ are described by 4^n^ + 2M, where *tf* denotes an n-gram term frequency vector (i.e n=2 in bigram and n=3 in trigram), and M denotes the number of chiralities. The total features of FV_2_ are described by M + 2N, where *cf* denotes SWCNT chirality features, N denotes the number of sequences and M denotes the number of chiralities. *a* is an indicator function for the analyte presence (either 0 or 1). The subscripts C, H, and Y represent CA-125, HE4, and YKL-40, respectively. **(B)** Each feature vector is processed by a multi-label classifier (black box) in order to classify (detect) each biomarker. IR is the intensity ratio and defined as IR=I/I_0_.

The second feature vector (FV_2_) uses chiralities as the example IDs (Fig. 3A_(2)_) combined with sequence-dependent optical responses as features. Underlying the FV_2_ is the hypothesis that a single SWCNT in combination with a number of DNA sequences is sufficient to determine the presence or concentration of biomarkers. SWCNTs were represented using the one-hot encoding (‘1’ for specific chirality and ‘0’ for the other chiralities)^54^ hence, the total number of features used for FV_2_ is 34 (12+2*11).

Input data formatted according to FV_1_ and FV_2_ were used to train several classification algorithms for the detection of individual biomarkers or combinations thereof (Fig. 3B). Three ML algorithms, support vector machine (SVM), random forest (RF), and artificial neural network (ANN), were trained using an initial dataset and were evaluated by 10-fold cross-validation. Bayesian optimization was used for hyperparameter tuning. The resulting F1-scores were used to assess model performance (Fig. S3).

### Classification model training and validation

We investigated the potential for the platform to detect the presence/absence of a single biomarker, HE4, using binary classification algorithms. We introduced the DNA-SWCNT complexes to solutions of HE4 and background/interferents FBS, BSA, and mixtures, all in PBS. We classified the data using several approaches such as bi-class (+/−HE4), multi-class (HE4, HE4+/−FBS, FBS, HE4+/−BSA, BSA), and multi-label (+/−HE4 and +/−FBS and +/−BSA). Criteria for excluding certain feature vectors were wavelength shifts higher than 20 nm or poor peak fitting, both most likely caused by low signal intensities. We found that RF resulted in better F1-scores than ANN and SVM (> 0.93) (Fig. S4A). The performance of bi-class classifiers was slightly better than multi-class and multi-label classifiers. Overall, the algorithms provided high F1-scores (>0.92). While using FV_2_, all algorithms provided high F1-scores (1.0 for bi-class and 0.9-1.0 for multi-class/multi-label classification). The high values of F1-scores raised concerns with overfitting, which could occur with small sample sets. Another concern is the high initial concentration of analytes in the training sets.

To alleviate those concerns and determine the detection limit of the platform for HE4 classification using the model trained with high concentrations, we tested against several lower HE4 concentrations. Fig. 4A shows F1-scores for the three algorithms using both FVs for 10 and 50 nM HE4 thresholds (as these concentrations are relevant to the cancer diagnosis). Both FVs generated high values for F1-scores on cross-validation at 50 nM HE4 concentration (F1-score > 0.89 in FV_1_ and F1-score > 0.98 in FV_2_) and in the range of 0.79-0.85 at 10 nM HE4. While both FVs continued to predict with high F1-scores, the performance of FV_2_ was better than FV_1_ and provided good results for all three algorithms.

**Figure 4.**
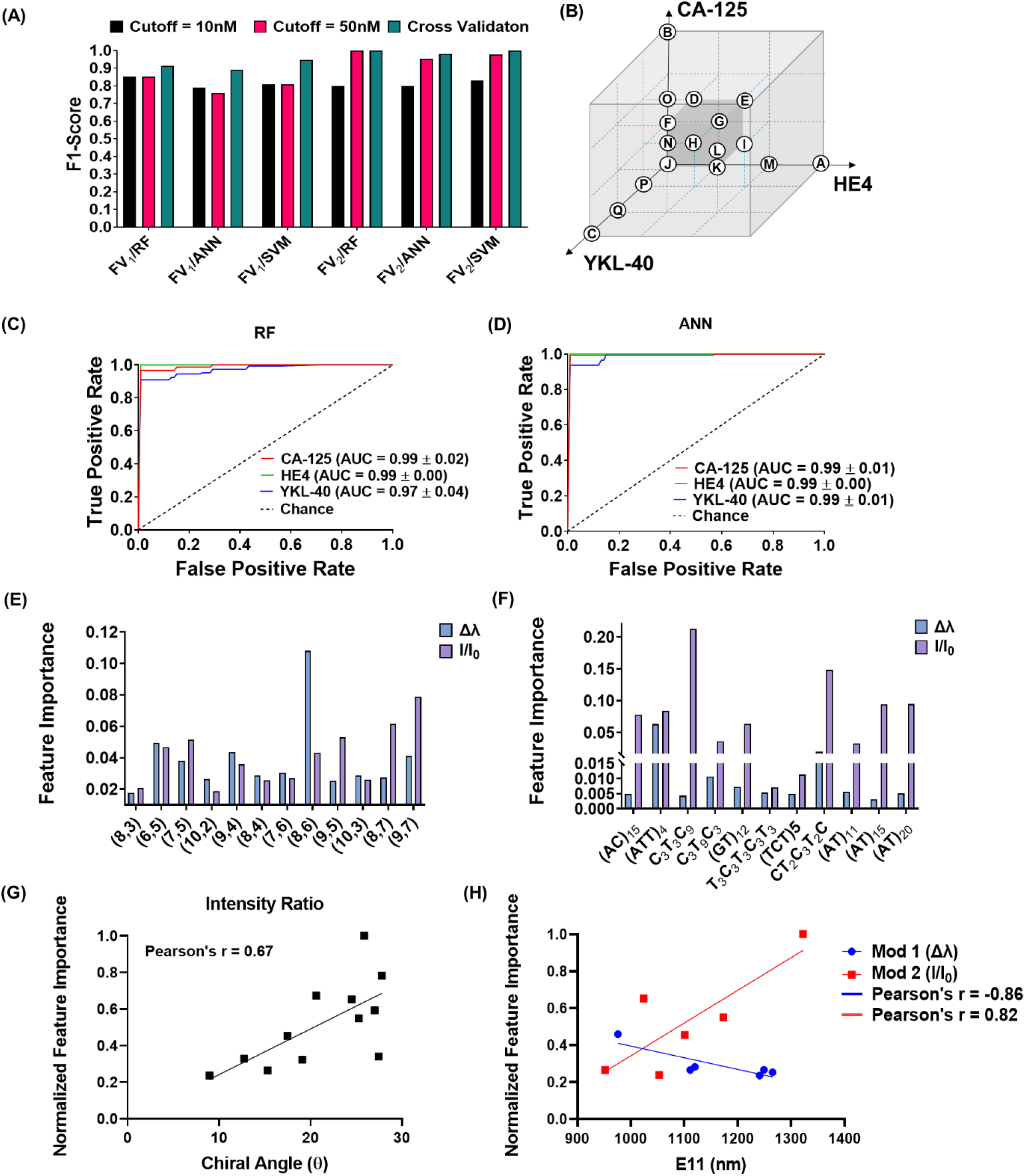
Machine-Perception Nanosensor Results and Analysis. **(A)** F1-scores of three algorithms for each FV corresponding to different thresholds. **(B)** Schematic of training/testing method for different concentrations of biomarker combinations. **(C)** ROC of each biomarker via RF model. AUC values for each biomarker. **(D)** ROC of each biomarker via ANN model. AUC values for each biomarker. **(E)** Feature importance of SWCNT chiralities generated by FV_1_. **(F)** Feature importance of DNA sequences generated by FV_2_. **(G)** Intensity change feature importance vs SWCNT chiral angle. **(H)** Normalized feature importance values of wavelength shift (Δλ) and intensity change vs. SWCNTs emission wavelength.

To investigate the potential for the platform to detect other/multiple cancer biomarkers, we optimized multilabel classification methodologies. We trained the ML algorithms using the optical response of the DNA-SWCNT complexes to a single and multiple combinations of HE4, CA-125, and YKL-40 with various concentrations, ranging from 0 nM to 100 nM (Fig. 4B, Table S1). In order to generate as comprehensive a dataset as possible, we screened over 17 different biomarker combinations, which resulted in more than 200 examples for each FV. We incubated the DNA-SWCNT complexes in FBS and PBS to assess the biomarkers in complex environments. We constructed three types of multi-label ML models: adaptive algorithm (AA), binary relevance (BR), and label powerset (LP) ^55,56^. Cross-validation results (Fig. S4B) show that the F1-scores using FV_2_ were significantly higher (>0.96) compared to FV_1_ (>0.68) across all the models, with RF and ANN outperforming SVM (with F1-scores of 0.97). To validate the F1-scores of the top-performing algorithms, we generated receiver operating characteristic (ROC) curves for the three biomarkers (Fig. 4C, D). The areas under the curve (AUC) were all greater than 0.97. Individual analyses of each biomarker showed high F1-scores for HE4 (1 and 0.99 in RF and ANN respectively), CA-125 (1 and 0.91 in RF and ANN respectively), and YKL40 (0.96 and 0.84 in RF and ANN respectively). These results demonstrate the ability of the model to detect single and multiple biomarkers in mixtures with high precision (Fig. S4C). In addition, the accuracy of detection was mostly high, depending on the biomarker (Fig. S4D).

Based on these results, we decided to proceed with FV_2_ as the feature vector used for the classification and RF and ANN as the ML algorithms. The better performances of FV_2_ suggest that the collection of spectroscopic features from a single SWCNT, in combination with a number of DNA sequences, is better than a feature vector that comprises data from a single DNA sequence on a number of SWCNTs.

To evaluate the concentration of each biomarker in each sample, we also conducted regression analysis. Regression results of RF and ANN using FV_2_ achieved R^2^ values of 0.93 and 0.92 respectively (shown in Fig. S4E).

### ML feature importance

To understand the relationship between the nanosensor array composition and the ML predictions, we used feature importance analysis to investigate the DNA-SWCNT properties that influence the prediction. We extracted the feature importance values from the algorithms using both FV_1_ and FV_2_ (Fig. 4E, F). We found that the relative importance of nanotube chiralities on the marker prediction appeared to correlate with chiral angle of the nanotube species, as defined by Pearson’s correlation coefficient (Fig. 4G). There also appeared to be some dependence on nanotube mod (Fig. S5 A,B), wherein nanotube chirality vectors (n,m) calculated via *mod*(n-m, 3) gives a value of 1 or 2 for semiconducting carbon nanotubes.^57,58^ We also found some correlation between the importances of wavelength shifting responses of mod 1 chiralities with the nanotube optical bandgap (E_11_) (r = −0.86) and intensity responses of mod 2 chiralities with optical bandgap (r = 0.82) (Fig. 4H, Fig. S5C-F). These correlations suggest that nanotube structure contributed to the differences in the optical responses of the nanosensors that enabled enough response diversity to result in positive predictive value.

Among DNA wrapping sequences, C_3_T_3_C_9_ and CT_2_C_3_T_2_C presented the highest and second highest feature importance values, respectively (Fig. 4F). Interestingly, the intensity ratio feature exhibited higher importance values than the wavelength shifting responses across all sequences. Using this feature importance analysis, we narrowed down the array to the five most important DNA sequences ((AC)_15_, (AT)_11_, (AT)_15_, CT_2_C_3_T_2_C,and T_3_C_3_T_3_C_3_T_3_) to reduce the number of features, and, therefore, the number of experimental conditions. The optimized model generated F1-scores of 0.98 for classification and R^2^ of 0.78 for regression. The combined results suggest that the sensitivity of this platform for the biomarkers is dependent on both the nanotube structure and the unique morphology of the DNA adhesion on the nanotubes due to sequence-dependent π-π stacking of the base pairs on the graphitic sidewall of the SWCNTs.

### Uterine lavage patient samples

Uterine washing samples were collected from consenting cancer patients with diagnoses of several gynecologic conditions, including ovarian and endometrial cancers (Fig. S6).^30,59–61^ To investigate the ability of the platform to detect multiple biomarkers in a patient biofluid sample, we tested the optimized MP platform in uterine washings. We incubated the DNA-SWCNT complexes in uterine lavage samples (N=22). The conventional clinical laboratory measurements showed a high biomarker distribution (Fig. 5A), with mean concentration values (in nanomolar) of HE4, CA-125, and YKL-40 equaling 2.75 ±0.63, 3.62 ±1.52, and 0.15 ±0.08, respectively. Due to the sub-nanomolar range of the biomarkers in the patient samples, we retrained the algorithms with lower concentrations of all biomarkers (1 pM-100 nM) and used the sensor array responses from the uterine lavage patient samples as a test set (Fig. 5B). The F1-score of the training set was improved as the threshold was decreased below 100 pM (0.95 increased to 0.97). In addition, the F1-score of the test set was significantly improved (0.93 increased to 0.99). This result indicates that several sample concentrations below 100 pM were inaccurately classified with the higher threshold. It is interesting to note that there was a negligible difference between the F1-scores when using a 10 pM or 1 pM threshold. This may be due to the fact that there was only one measurement below 10 pM. We further evaluated the F1-score of individual biomarker predictions (Fig. 5C). While there was an improvement in the F1-scores of each biomarker when the threshold was decreased to 10 pM (> 0.95), there were much more significant improvements in the sensitivities to CA-125 and YKL-40 (0.87 increased to 1 and 0.89 increased to 0.95 in CA-125 and YKL-40 respectively).

**Figure 5.**
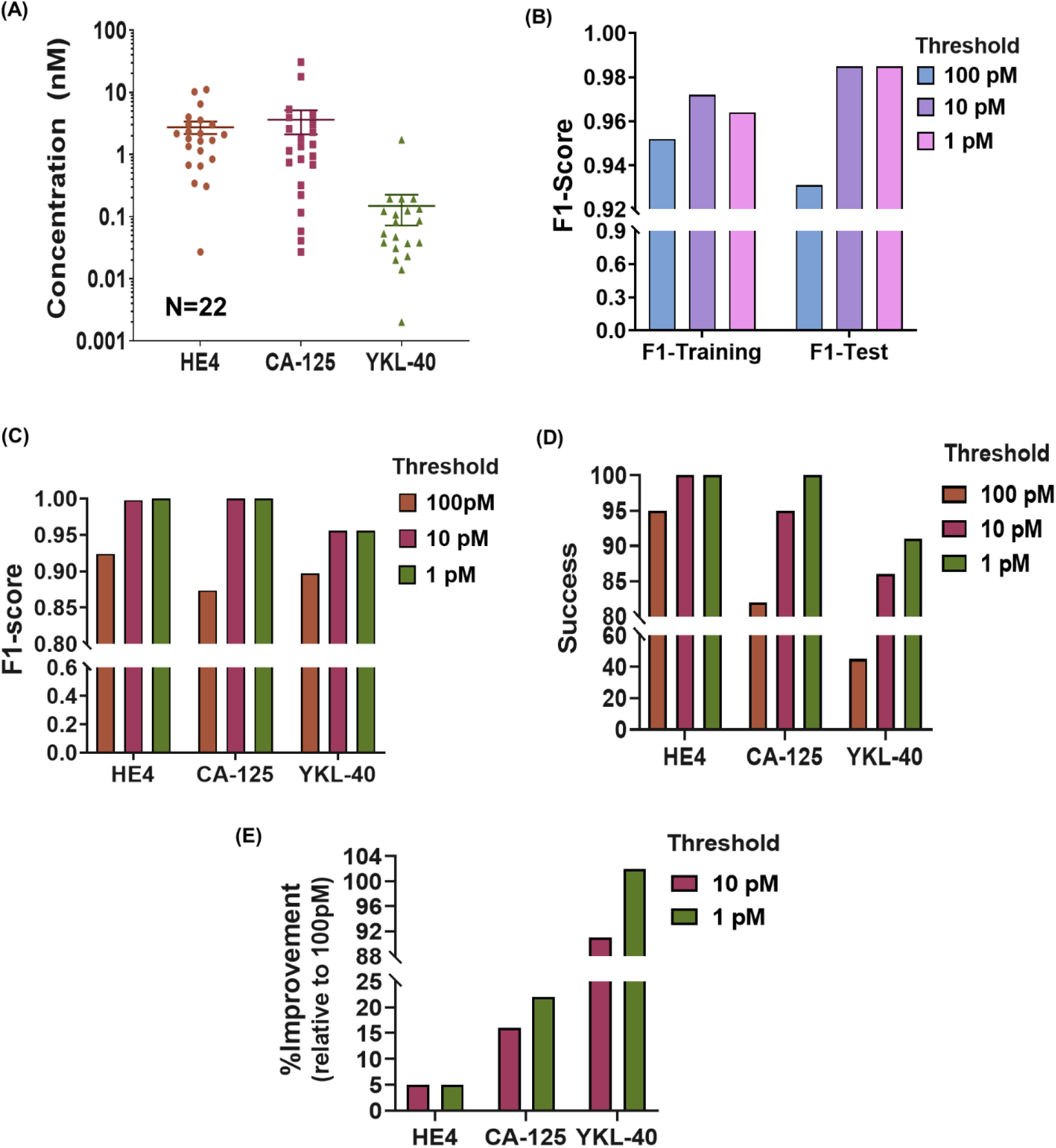
Biomarker detection in uterine lavage samples. **(A)** Concentrations of HE4, CA-125, and YKL-40 were measured by ELISA in uterine lavage samples. **(B)** Classification F1-scores for detection of the three biomarkers in lavage samples from training and test data sets, applying different protein concentration thresholds. **(C)** Classification F1-scores of nanosensor detection of each biomarker, applying different protein concentration thresholds. **(D)** The success of the detection of each biomarker via classification, applying different concentration thresholds. **(E)** Improvement in classification success, relative to the threshold of 100 nM.

To evaluate the prediction strength, we examined the success of classification results for each threshold by comparing them to the actual levels of each biomarker measured by the clinical laboratory (Fig. 5D). We defined success as the percentage of correct classification (either true positive or true negative) for each biomarker. In all the biomarkers, the classification prediction was significantly improved when the threshold was decreased below 100 pM (Fig. 5E). HE4 presented the most successful classifications and showed improvement from 95% to 100% success with both 10 pM and 1 pM thresholds. CA-125 was initially predicted with 82% success but significantly improved to 100% success when the threshold was changed to 1 pM. The most significant change was observed with YKL-40, from 50% success with a 100 pM threshold to 91% success with a 1 pM threshold. The machine-perception platform was able to accurately classify the patient biofluid samples, indicated by the high F1-scores and classification success values.

## Discussion

Current diagnostic methodologies are one-to-one recognition assays that mainly use antibodies. Herein, we described a new approach for the detection of multiple biomarkers in biofluids for disease diagnosis using an artificial molecular perception system. We developed an array of relatively non-specific DNA-SWCNT sensors, containing individual hybrids of 132 DNA-wrapped SWCNTs. The use of multiple SWCNT chiralities enabled us to generate a large set of sensors that could be interrogated rapidly via high-throughput near-infrared spectroscopy to form a wide diversity of responses when they were exposed to different target proteins. Based on several studies,^34,62–64^ we initially targeted gynecologic cancer biomarkers HE4, CA-125, and YKL-40.

Advantages of the method include the high optical sensitivity of SWCNTs to diverse analytes and the ability to modify their environmental sensitivities/specificities. We introduced a diverse set of SWCNT environmental responsivities via surface coatings of different DNA sequences which modulated the optical bandgaps and surface morphologies. Machine learning algorithms enabled training from DNA-SWCNT spectral response data to detect biomarkers in both lab-generated samples and cancer patient uterine lavage samples.

Notably, the classification success rate in patient samples was high even in sub-nanomolar ranges, with a rate of 100% for HE4 and CA-125 and 91% in YKL-40. These results support the conclusion that the perception mode of sensing can successfully generate accurate predictions. Combining detection and quantification will allow this technology to better screen and categorize patients based on the levels of markers for early detection.

This platform could be continuously improved by increasing the sizes of datasets and analyzing feature importance. For example, interpreting the feature importance values can aid with DNA sequence design and expanding the library of DNA-SWCNT complexes. Also, expanding the spectroscopic range may increase the number of SWCNT chiralities, and thus, sensors that can be measured. While increasing the number of features (nanosensors) may contribute to the sensitivity of the platform, the number of examples (conditions) should be increased as well (to prevent overfitting). We also recognize the need to increase the number of patient samples to continually validate and increase the robustness of the model.

Finally, due to the flexibility and the non-specific nature of the individual sensor elements, the proposed MP platform is not restricted to ovarian cancer biomarkers and can potentially be trained to detect other disease biomarkers without the need to engineer different arrays of nanosensors. This platform enables antibody-free detection that would be useful when an especially robust or long-term measurement is needed, such as in wearable/implantable devices, point-of-care diagnostics, and for under-resourced situations where cold chain storage may not be available.

## Materials and Methods

### Materials

Single-walled carbon nanotubes (SWCNTs) produced by the HiPco process were purchased from Unidym (Sunnyvale, CA, USA). Single-stranded DNA (ssDNA) oligonucleotides (T_3_C_3_T_3_C_3_T_3_; C_3_T_9_C_3_; C_3_T_3_C_9_; (TCT)_5_; (GT)_12_; (AT)_11_; (AT)_15_; (AT)_20_; (ATT)_4_; (AC)_15_ and CT_2_C_3_T_2_C) were purchased from IDT DNA (Coralville, IA, USA). Human epididymis protein 4 (HE4) was purchased from RayBiotech. Human cancer antigen 125 (CA-125), also known as MUC16, and Human chitinase 3-like 1 (YKL-40) were purchased from R&D Systems. Bovine serum albumin was purchased from Sigma Aldrich. Fetal bovine serum was purchased from Thermo Fisher Scientific. Uterine washings from IRB consented cancer patients were provided by the Department of Laboratory Medicine at Memorial Sloan Kettering.

### Preparation of DNA-Wrapped Single-Walled Carbon Nanotube Complexes

Single-walled carbon nanotubes (SWCNTs) were mixed with a specific DNA oligonucleotide at a 1:2 mass ratio, respectively, in 1 mL of IDTE buffer. The sample was ultrasonicated continuously for 45 minutes at 40% of maximum amplitude, using a 3 mm titanium tip (SONICS Vibra Cell). The mixture was ultracentrifuged (Sorvall Discovery 90SE) for 30 minutes at 250,000xg. The top 80% of the supernatant was collected and the concentration of suspended SWCNTs was determined by UV/Vis/NIR spectrophotometry (JASCO V-670) using the extinction coefficient A_910_ = 0.02554l mg^-1^cm^-1^; where the path length l is 1 cm. In order to remove excess free DNA, 300 µL of the sample was filtered twice using a 100 kDa Amicon centrifuge filter (Millipore) at 5,000xg for 10 minutes. Following filtration, the DNA-SWCNT complexes were tested at a concentration of 5 mg/L SWCNT in 100 µL solution in 96 well plates. Zeta potential of the DNA-SWCNT complexes was measured using a Zetasizer ZSP (Malvern). The samples were diluted to a concentration of 0.5mg/L using double distilled water.

### Near-Infrared Fluorescence Spectroscopy of DNA-SWCNTs

Near-infrared fluorescence spectroscopy was used to measure the photoluminescence emission from the DNA-SWCNT complexes, as described previously.^65^ For solution measurements, spectra were acquired using an apparatus built in-house consisting of a continuous wave 730 nm diode laser with an output power of 2 W or a SuperK EXTREME supercontinuum white-light laser source connected to a Varia variable bandpass filter accessory capable of tuning the output from 490–825 nm with a bandwidth of 20 nm (NKT Photonics). The laser was injected into a multimode fiber that was fed into the back of an Olympus IX-71 inverted microscope where it passed through a 20x LCPlan N, 20x/0.45 objective (Olympus, USA) and a dichroic mirror (875 nm cut-off; Semrock). The light was f-number matched to the spectrometer using several lenses and injected into an IsoPlane spectrograph (Princeton Instruments) with a slit width of 410 µm which dispersed the emission using a 86 g mm-1 gating with a 950 nm blaze wavelength coupled to a NIRvana 2D InGaAs near-infrared detector (Princeton Instruments) or a Shamrock 303 Spectrometer with Andor iDus 1D InGaAs Array Camera (Oxford Instruments). An HL-3-CAL EXT halogen calibration light source (Ocean Optics) was used to correct for wavelength-dependent features in the emission intensity arising from the excitation power, spectrometer, detector, and other optics. A Hg/Ne pencil-like calibration lamp (Newport) was used to calibrate spectrometer wavelengths. Data were obtained from each well of a 96-well plate using the custom LabVIEW (National Instruments) code. Another custom program, written in MATLAB (MathWorks) software was used to subtract background, correct for abnormalities in excitation profiles, and fit the data with Lorentzian functions. Smoothing, where applicable, was done by applying a Savitzky-Golay filter.

### Atomic Force Microscopy

DNA-SWCNT complexes were plated on a freshly cleaved mica substrate (SPI) for 4 minutes before washing with 10 ml of distilled water and blowing dry with argon gas. An Asylum Research MFP-3D-Bio instrument equipped with an Olympus AC240TS AFM probe in alternating-current mode was used. Data were acquired at 2.93 nm pixel −1 x-y resolution and 15.63 pm z resolution. The images were analyzed using Gwyddion software. To measure height or length distributions, at least 20 ROIs were analyzed.

### Machine Learning Method Development

The dataset comprises the photoluminescence spectra of each combination of DNA-SWCNT complex exposed to different combinations of a small number of analytes (HE4, CA-125, YKL-40, BSA, FBS). That is, we had total N·M·L combinations where N is the number of DNA sequences, M is the number of SWCNT chiralities, and L is the number of analyte combinations. The spectra were analyzed to yield two parameters for each SWCNT type: the wavelength peak shift (Δ*λ*) and intensity ratio (IR):

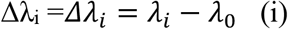

and

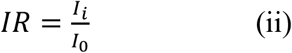

where *λ*_0_ and *I*_0_ are the wavelength and intensity of a control sample (DNA-SWCNT without analyte); *λ*_*i*_and *I*_*i*_ are the wavelength and intensity of DNA-SWCNT with analyte combination, *i*. Input and output (target) variables were identified for the machine learning algorithms. The input variables include DNA sequence, SWCNT chirality, and the two spectroscopically measured parameters (Δ*λ*_*i*_, IR). The output variable either represents the presence (for classification) or concentration (for regression) of each analyte. Three classification approaches were examined: bi-class (+/− biomarker), multi-class (+/ biomarker combination), and multi-label (+/− each biomarker).

To train the models, categorical data (such as SWCNT chirality and analyte type) were transformed to numeric values, using the one-hot encoding technique ^54^. DNA sequences were encoded as term-frequency vectors, using subsets of two or three bases as a term and calculating the frequency of that term in the sequence.^47^ Fig. S3 depicts the overall scheme for the input feature construction. Two feature vectors were constructed to emphasize the sensitivity of each component in the DNA/SWCNT complex and find a balance between the number of features and examples. For each feature vector, one component of the DNA/SWCNT complex was encoded as an ID of the example while the other components’ responses were defined as features. In feature vector 1 (FV_1_), the DNA sequences were encoded as the IDs and the chirality-dependent optical responses as features. In feature vector 2 (FV_2_) the SWCNT chiralities were encoded as the IDs and the sequence-dependent optical responses as features.

Three algorithms, support vector machine (SVM), random forest (RF), and artificial neural network (ANN) were trained and tested with FV_1_ and FV_2_ for both classification and regression. Each model was evaluated by 10-fold cross-validation. All machine learning algorithms were implemented using the Scikit-learn machine learning library.^55^ In order to find hyperparameters that maximize performance, Bayesian hyperparameter optimization was implemented using HyperOpt.^66^

Each model was evaluated by the produced F1-score and accuracy values for classification and R^2^ value for regression. Accuracy (Eqn. S1) calculates the percentage of correctly classified examples. F1-score, which is a composite value of precision (Eqn. S2) and recall (Eqn. S3), gives a measure of accuracy but takes the false positives and negatives into account as well (Eqn. S4).

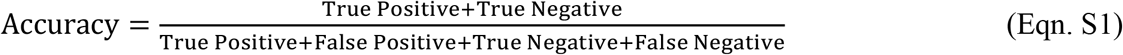

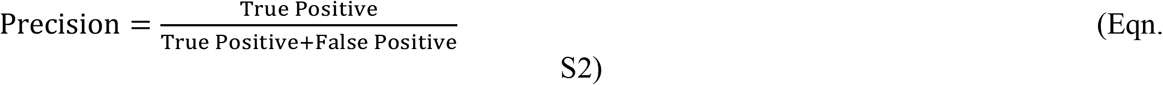

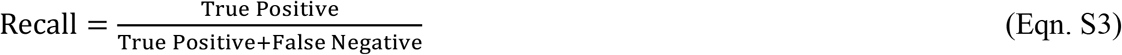

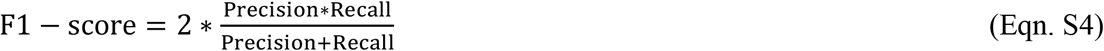

### CA-125 Concentration Unit Conversion

The concentration unit of CA-125 used in the clinic, Units/ml, was converted to nanomolar by titrating CA-125 and measuring using an ARC i2000 instrument. A linear concentration curve with R^2^=0.9997 was generated. The resulting unit conversion is as follows: [Unit/ml] = 0.18*[pM].

### Statistical Analysis

*In vitro* experiments were analyzed by two-sided t-tests. Reported p-values were assigned **** = p < 0.0001, *** = p < 0.001, ** = p < 0.01, * = p < 0.05 and exact p-values are reported in captions.

## Acknowledgments

This work was supported in part to DAH by the NIH New Innovator Award (DP2-HD075698), NCI (R01-CA215719), the Cancer Center Support Grant (P30 CA008748), the National Science Foundation CAREER Award (1752506), the American Cancer Society Research Scholar Grant (GC230452), the Honorable Tina Brozman Foundation for Ovarian Cancer Research, the Ara Parseghian Medical Research Fund, the Expect Miracles Foundation - Financial Services Against Cancer, the Pershing Square Sohn Cancer Research Alliance, the Cycle for Survival’s Equinox Innovation Award in Rare Cancers, Mr. William H. Goodwin and Mrs. Alice Goodwin, the Kelly Auletta Fund and the Commonwealth Foundation for Cancer Research, the Experimental Therapeutics Center, and the Center for Molecular Imaging and Nanotechnology of Memorial Sloan Kettering Cancer Center; to ML by NIGMS (R35GM134878), Functional Genomic Initiative, the Alan and Sandra Gerry Metastasis and Tumor Ecosystems Center. Z.Y. was supported by the Ann Schreiber Mentored Investigator Award (Ovarian Cancer Research Fund) and Young Investigator 2019 (Kaleidoscope of Hope). The authors would like to thank the Molecular Cytology Core Facility at Memorial Sloan Kettering Cancer Center. We would like to thank Biran Wang for acquiring the DNA-SWCNT AFM images. We would like to thank Biorender.com for serving as a platform for scientific illustration. YY was supported by a Dean’s Fellowship at Lehigh University., AJ’s contributions are part of the NHI initiative at Lehigh University, M.Z. acknowledges NIST internal funding for support. DAL is supported by US Department of Defense Award: W81XWH-15-1-0429, The Honorable Tina Brozman Foundation for Ovarian Cancer Research, and Arnold Chavkin and Laura Chang.

## Supplementary Materials

Figure S1. Characterization of DNA-SWCNT complexes.

Figure S2. Optical response of DNA-SWCNT

Figure S3. Overall scheme for input feature construction.

Figure S4. F1-scores of MP models.

Figure S5. Feature Importance Analysis.

Figure S6. Patients’ diagnosis distribution.

Figure S7. Concentration unit conversion of CA-125.

Table S1. Data used to construct the plot in Figure 4B.

