## Supplementary Materials for "Machine-Perception Nanosensor Platform to Detect Cancer Biomarkers"

**This PDF file includes:**

Figure S1. Characterization of DNA-SWCNT complexes.

Figure S2. Optical response of DNA-SWCNT

Figure S3. Overall scheme for input feature construction.

Figure S4. F1-scores of MP models.

Figure S5. Feature Importance Analysis.

Figure S6. Patients’ diagnosis distribution.

Figure S7. Concentration unit conversion of CA-125.

Table S1. Data used to construct the plot in Figure 4B.

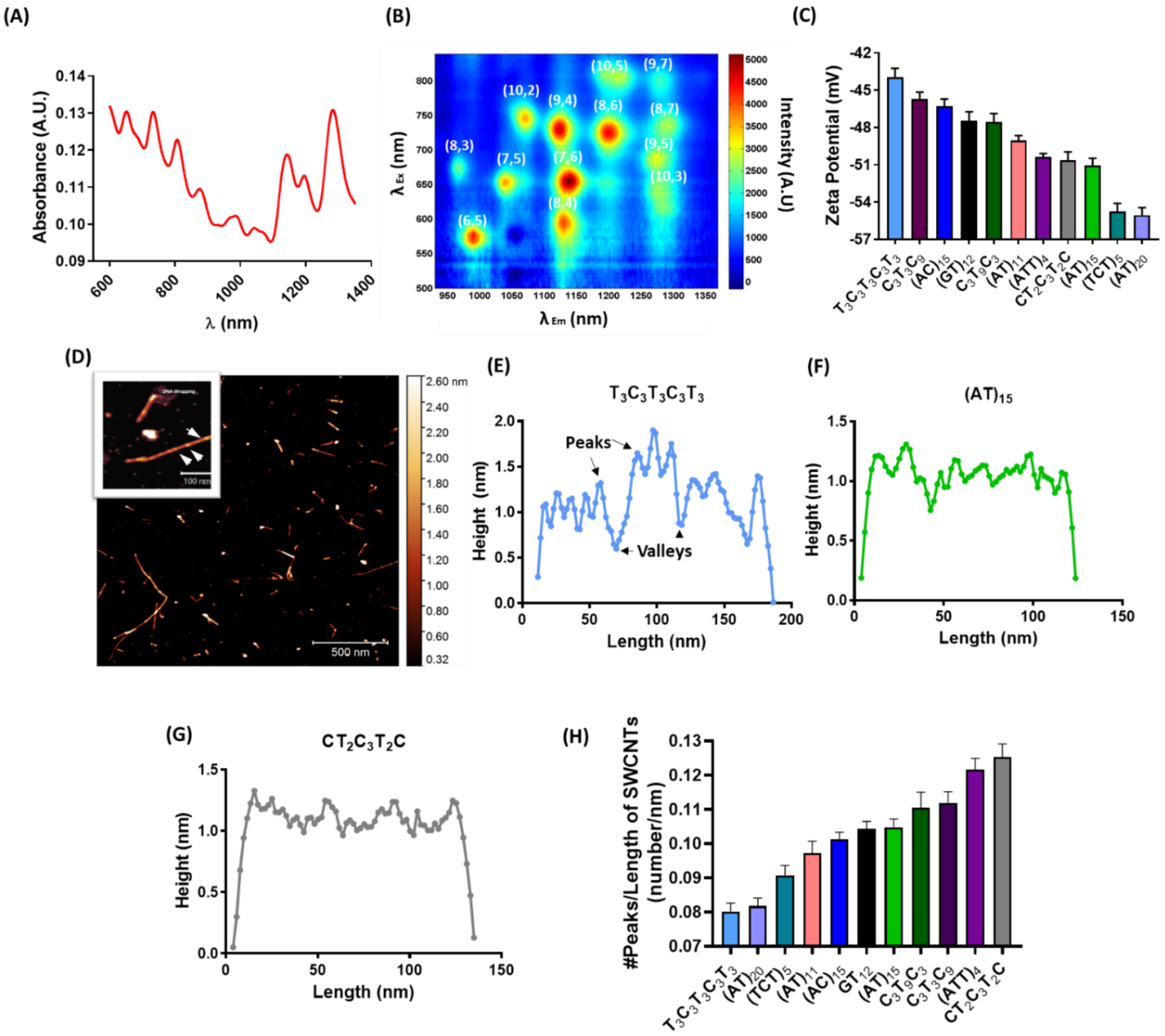


**Figure S1. Characterization of DNA-SWCNT complexes. (A)** Representative absorbance spectrum of a DNA-SWCNT complex, $(AC)_{15}$-SWCNT. **(B)** 3D photoluminescence plot of the $(AC)_{15}$-SWCNT complex. **(C)** Zeta potentials of the DNA-SWCNT complexes; data were averaged over 7 acquisitions. **(D)** AFM image of $(AC)_{15}$-SWCNTs. White arrows denote height maxima ascribed to ssDNA. **(E)** Height profile along the length of an individual $T_3C_3T_3C_3T_3$-SWCNT complex. **(F)** Height profile along the length of an individual $(AC)_{15}$-SWCNT complex. **(G)** Height profile along the length of an individual $CT_2C_3T_2C$-SWCNT complex. **(H)** The average density of DNA peaks per unit length on the DNA-SWCNT complexes, quantified from AFM images; data were averaged over 20 acquisitions.

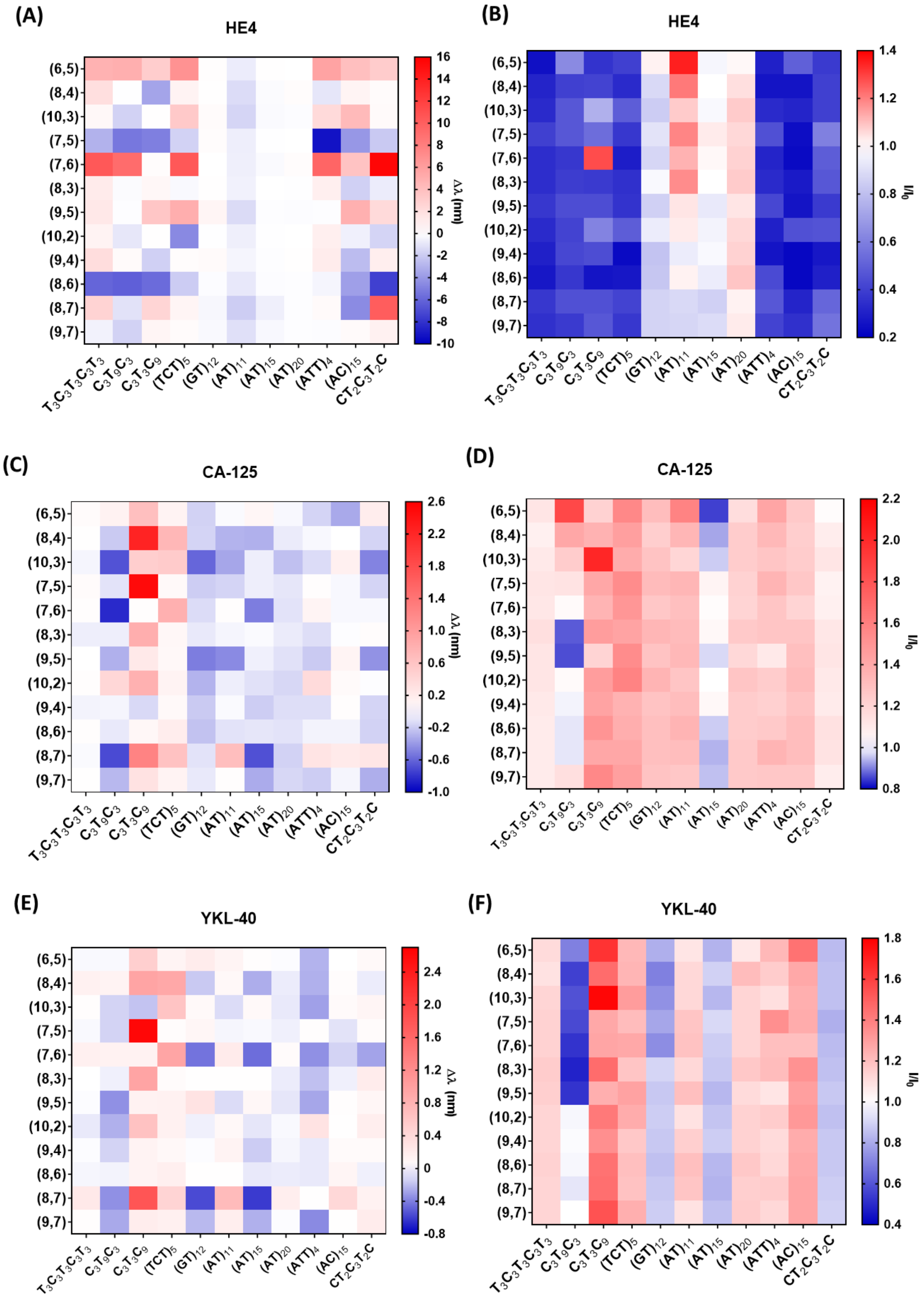

(A)
HE4
(6,5)
(8,4)
(10,3)
(7,5)
(7,6)
(8,3)
(9,5)
(10,2)
(9,4)
(8,6)
(8,7)
(9,7)
$T_3C_3T_3C_3T_3$
$C_3T_9C_3$
$C_3T_3C_9$
$(TCT)_5$
$(GT)_{12}$
$(AT)_{11}$
$(AT)_{15}$
$(AT)_{20}$
$(ATT)_4$
$(AC)_{15}$
$CT_2C_3T_2C$
Δλ (nm)
16
14
12
10
8
6
4
2
0
-2
-4
-6
-8
-10
(B)
HE4
$I/I_0$
1.4
1.2
1.0
0.8
0.6
0.4
0.2
(C)
CA-125
Δλ (nm)
2.6
2.2
1.8
1.4
1.0
0.6
0.2
-0.2
-0.6
-1.0
(D)
CA-125
$I/I_0$
2.2
2.0
1.8
1.6
1.4
1.2
1.0
0.8
(E)
YKL-40
Δλ (nm)
2.4
2.0
1.6
1.2
0.8
0.4
0
-0.4
-0.8
(F)
YKL-40
$I/I_0$
1.8
1.6
1.4
1.2
1.0
0.8
0.6
0.4

**Figure S2. Optical responses of DNA-SWCNT complexes to cancer biomarkers. (A)** Heatmap of wavelength shifting responses of DNA-SWCNT complexes upon incubation with 100 nM HE4 in 10 % FBS in PBS. **(B)** Heatmap of intensity changes of DNA-SWCNT complexes upon incubation with 100 nM HE4 in 10% FBS in PBS. **(C)** Heatmap of wavelength shifts of DNA-SWCNTs complexes upon incubation with 100nM CA-125 in 10 % FBS in PBS. **(D)** Heatmap of intensity changes of DNA-SWCNT complexes upon incubation with 100 nM CA-125 in 10% FBS in PBS. **(E)** Heatmap of wavelength shifts of DNA-SWCNTs complexes upon incubation with 100 nM YKL-40 in 10% FBS in PBS. **(F)** Heatmap of intensity changes of DNA-SWCNTs complexes upon incubation with 100 nM YKL-40 in 10 % FBS in PBS.

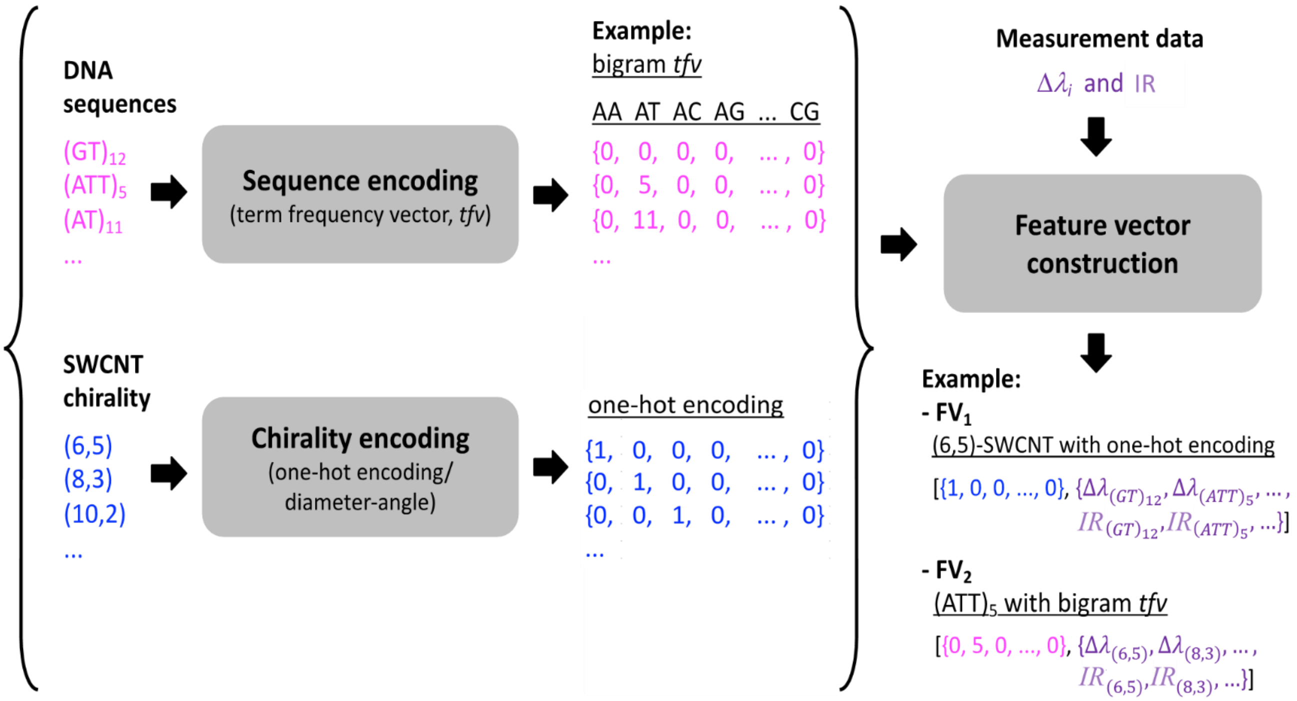


**Figure S3. Scheme for input feature construction.** The DNA sequence was encoded to a numeric vector using the term-frequency method (*47*) and SWCNT chirality was transformed into numeric vectors using a one-hot encoding technique. The numeric vectors could be then combined with a set of measurement data ($\Delta\lambda_i$, IR) using the $FV_1$ and $FV_2$ construction methods. This figure presents an example of a feature vector for a single example obtained by the $FV_1$ and $FV_2$ methods.

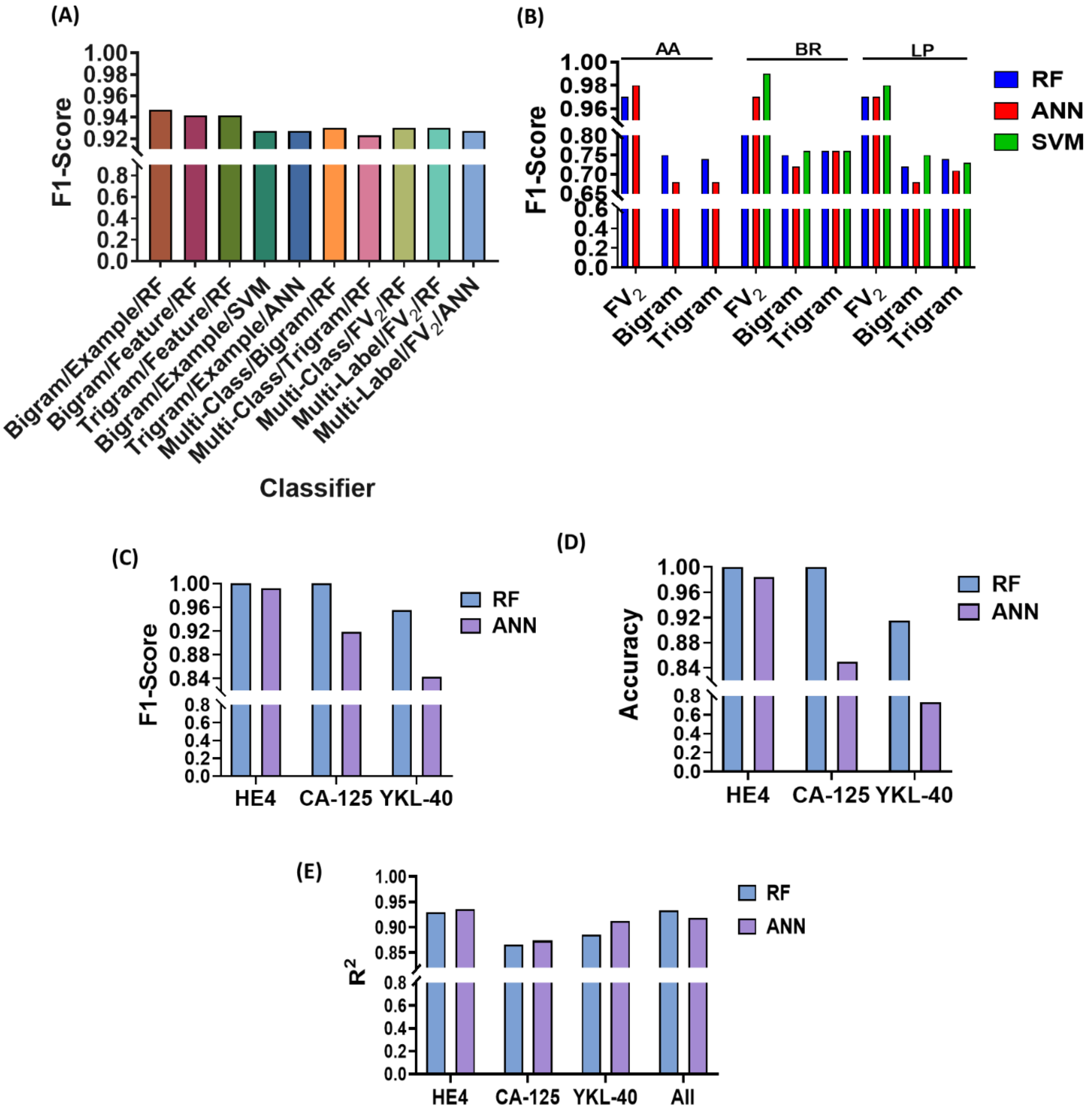


**Figure S4. Comparison of machine learning models. (A)** F1-scores of different ML algorithms and FV sets. 'Example' denotes complete data of examples after removing incomplete sets of specific examples. 'Feature' denotes the complete data set of features after removing outliers results of specific features. 'Multi-Label' denotes the classification of single biomarkers in a mixture. 'Multi-Class' denotes the classification of single or mixed biomarkers with other analytes. **(B)** F1-scores of different ML algorithms and FV sets using different multi-label classifiers such as adaptive algorithm (AA), binary relevance (BR), and label powerset (LP). **(C)** F1-scores of each biomarker classification using random forest (RF) and artificial neural network (ANN) models. **(D)** Accuracy of each biomarker classification using RF and ANN models. **(E)** $R^2$ values from regression analysis denoting the accuracy of quantitative prediction of the concentration of each biomarker, as well as the combined accuracy.

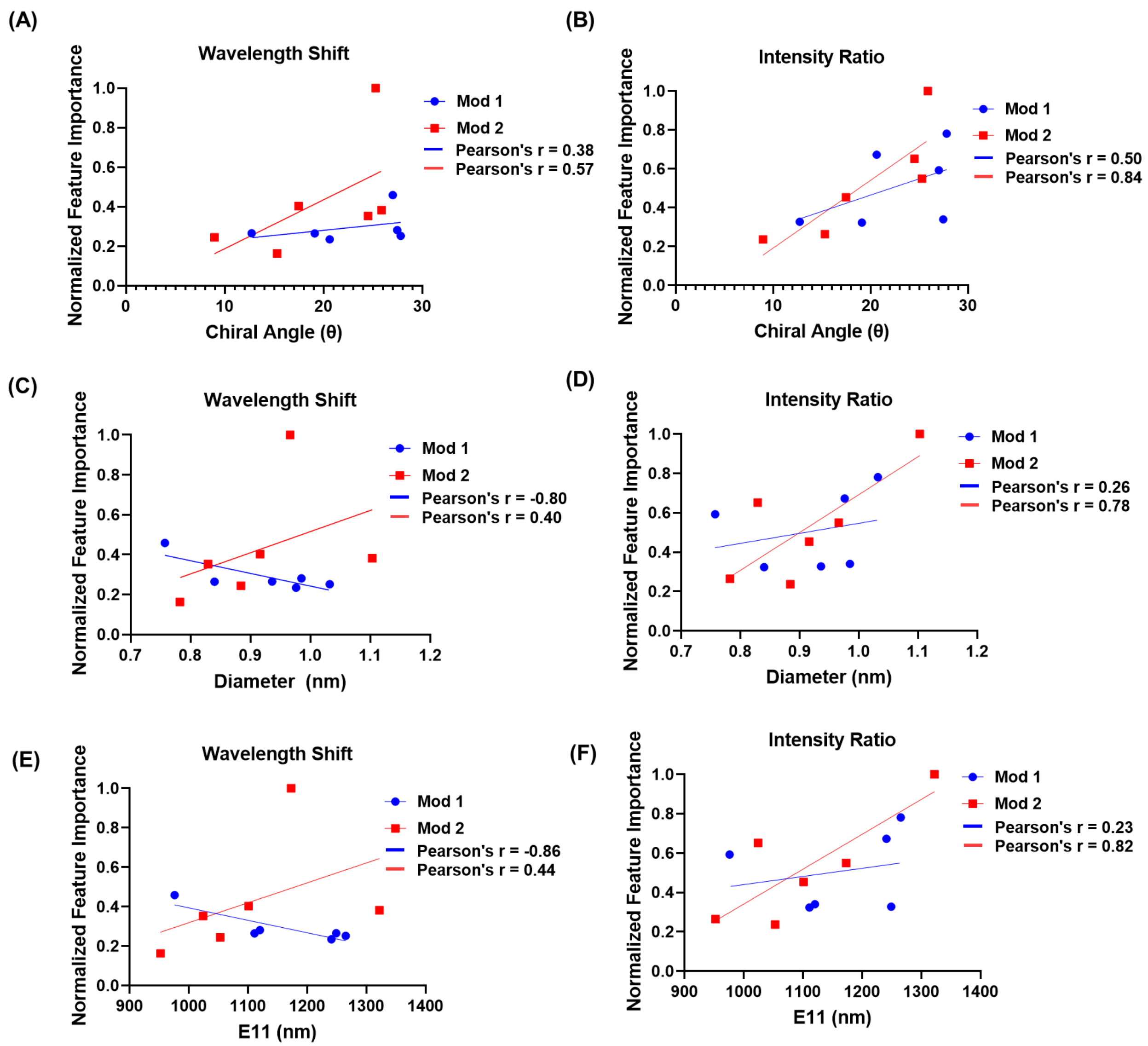


**Figure S5. Feature Importance Analysis. (A)** Normalized feature importance values of wavelength shift and **(B)** intensity change of 12 SWCNT species/chiralities, plotted by nanotube chiral angle. **(C)** Normalized feature importance values of wavelength shift and **(D)** intensity change of 12 SWCNT species/chiralities, plotted by nanotube diameter. **(E)** Normalized feature importance values of wavelength shift and **(F)** intensity change of 12 SWCNT species/chiralities, plotted by nanotube emission wavelength.

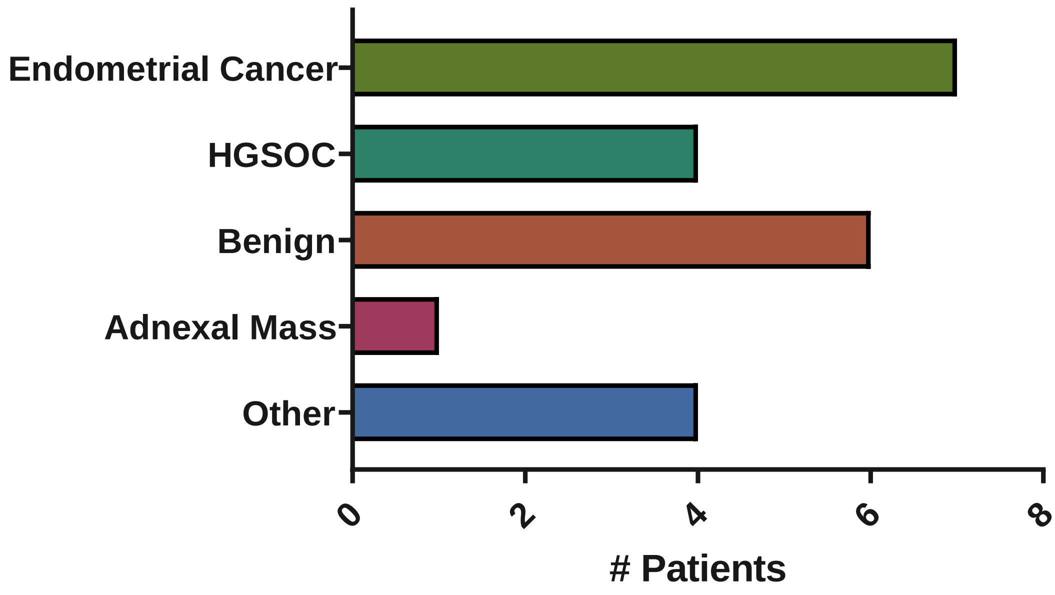


**Figure S6. Uterine lavage sample patient diagnoses.** Diagnoses of patients providing uterine lavage samples in this study; HGSOC = high grade serous ovarian carcinoma.

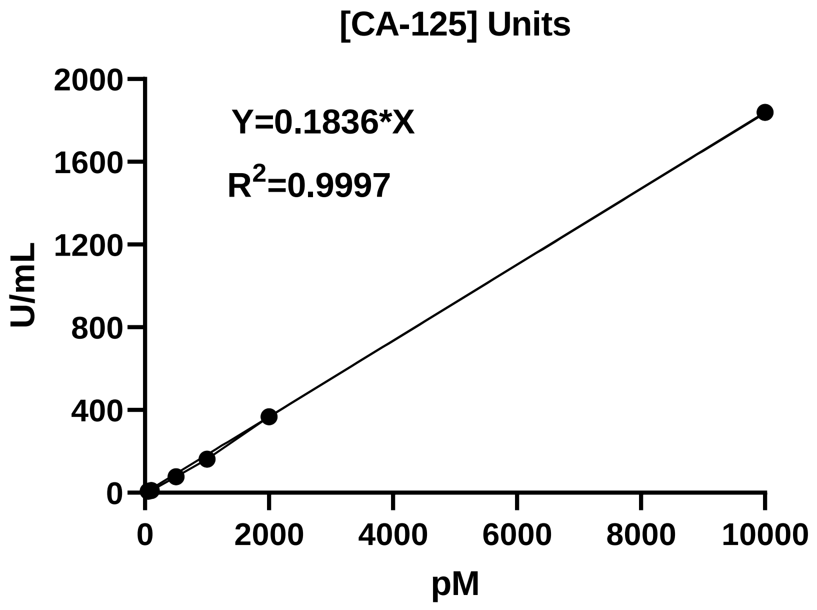


**Figure S7. Concentration unit conversion of CA-125.** The linear curve of Units/mL vs. concentration in pM.

**Table S1. Data used to construct the plot in Figure 4B.**

| | Concentration (nM) | | |
|---|---|---|---|
| | **HE4** | **CA-125** | **YKL-40** |
| **A** | 100 | 0 | 0 |
| **B** | 0 | 100 | 0 |
| **C** | 0 | 0 | 100 |
| **D** | 30 | 70 | 30 |
| **E** | 70 | 70 | 30 |
| **F** | 30 | 70 | 70 |
| **G** | 70 | 70 | 70 |
| **H** | 30 | 30 | 30 |
| **I** | 70 | 30 | 30 |
| **J** | 30 | 30 | 70 |
| **K** | 70 | 30 | 70 |
| **L** | 30 | 0 | 0 |
| **M** | 70 | 0 | 0 |
| **N** | 0 | 30 | 0 |
| **O** | 0 | 70 | 0 |
| **P** | 0 | 0 | 30 |
| **Q** | 0 | 0 | 70 |
